# Evaluating the quality of brainstem ROI registration using structural and diffusion MRI

**DOI:** 10.1101/2025.09.22.675000

**Authors:** Yi-An A. Chen, Lars Kasper, Clement T. Chow, Yu Kuo, Alexandre Boutet, Jürgen Germann, Andres M. Lozano, Kâmil Uludağ, Andreea O. Diaconescu, Sriranga Kashyap

## Abstract

Accurate registration of regions of interest (ROIs) from standard atlases to participants’ native spaces is a critical step in fMRI studies, as it directly affects the reliability of sampled BOLD signals. While T1-weighted (T1w) image-based ROI registration is well validated and widely adopted in cortical fMRI, its performance degrades in brainstem studies due to the small size, dense packing, and poor visibility of brainstem nuclei on T1w contrast. We hypothesized that incorporating diffusion MR images, containing more information about internal brainstem architecture, should improve ROI registration accuracy. To test this, we developed four registration pipelines that either included or excluded diffusion-based alignment components and evaluated their performance using data from n=20 healthy participants. Registration accuracy was assessed using Dice coefficient for the red nucleus (RN) and the substantia nigra (SN), and mis-registration fraction—a metric developed for nuclei that cannot be manually delineated—for the dorsal raphe nucleus (DRN). The results showed that diffusion-based pipelines, using fractional anisotropy (FA) images, non-diffusion-weighted (b0) images, and multivariate combination, outperformed the T1w-only baseline. Probabilistic maps derived from inverse-transformed native ROIs further supported improved sensitivity to inter-individual anatomical variability in the diffusion-augmented pipelines. In addition, analysis of gradient magnitude maps from the Jacobian determinants revealed associations between localized deformation and image modality-specific landmarks. These findings demonstrate the potential of diffusion-augmented pipelines for improving brainstem ROI registration, which could enhance the robustness of fMRI studies on brainstem disorders characterized by functional dysregulation.

## 1 Introduction

The brainstem is a crucial hub of the central nervous system. It serves as a conduit for numerous ascending and descending axonal tracts, actively linking the brain with the rest of the body to coordinate physiological functions (Fernández-Gil et al., 2010). As one of the most evolutionarily conserved structures (Pessoa et al., 2019), the brainstem harbors several highly specialized, small neuronal clusters, known as brainstem nuclei, that support autonomic and neuromodulatory functions essential to vital bodily processes. Despite its importance, the brainstem has been relatively underexplored in functional MRI (fMRI) research due to challenges arising from a combination of its anatomical location and constraints from image acquisition. These include increased sensitivity to motion artifacts from pulsating surface arteries (e.g., basilar and vertebral arteries), signal dropouts and image distortion at air-tissue interfaces (e.g. sphenoid sinus, pharynx), and reduced signal-to-noise ratio (SNR) resulting from the small size of brainstem nuclei (Beissner, 2015; Sclocco et al., 2018). Collectively, these factors amplify the impact of co-registration errors between functional and anatomical data, compromising the validity and reliability of brainstem fMRI. To address these brainstem-specific limitations and enhance the feasibility of brainstem fMRI as a research tool, the fMRI community has been actively developing tailored data acquisition and preprocessing strategies to improve the fidelity of brainstem imaging (Beissner, 2015; Sclocco et al., 2018; Matt et al, 2019; Cauzzo et al., 2022; Singh et al., 2022; Hansen et al., 2024, Zhang et al., 2025).

In conventional fMRI analysis, cortical and large subcortical (e.g. basal ganglia, thalamus) ROIs, defined using anatomical data and atlases, are used to extract BOLD time series for statistical analysis. This process typically follows a two-step approach involving spatial transformation—that is, a mathematical mapping between coordinate systems—between the participant’s “native space,” defined by participant-specific anatomical data, and a “standard space,” defined by population-based brain templates (e.g., MNI152 or IIT templates) constructed by averaging data from dozens to hundreds of individuals (Fonov et al. 2009). In the first step, a nonlinear spatial transform from standard space to native space is calculated by optimizing alignment between a standard T1-weighted (T1w) image template and the participant’s high-resolution T1w image. In the second step, this transform is applied to map ROIs from an atlas defined in standard space into the participant’s native space. This T1w-based approach generally enables accurate registration of cortical and forebrain subcortical ROIs due to the excellent gray matter-white matter contrast provided by T1w images. As this method is considered standard practice and is implemented in the default preprocessing pipelines of many fMRI analysis toolboxes, it is also widely used in brainstem fMRI studies, except in rare cases where certain brainstem nuclei can be directly visualized in native space due to their distinct biochemical properties and where additional acquisition time is acceptable for the studied population (Chen et al., 2014; Priovoulos et al., 2018; Mazancieux et al., 2023). Unlike the cortex, however, the brainstem lacks well-defined macroscopic gray matter-white matter boundaries, with nuclei blending into a dense network of interwoven axonal tracts. Consequently, the brainstem often appears in T1w images as a relatively homogeneous structure, with clear external boundaries yet unclear internal architecture. As a result, when applied to brainstem fMRI, the first-step standard-to-native registration relies primarily on gross anatomical features. This can lead to suboptimal registration of brainstem ROIs because the spatial locations of brainstem nuclei are governed more by internal microstructural organization than by gross surface contours and there is known inter-individual variability of the internal organization (Shepherd et al., 2020; Bianciardi et al., 2018; García-Gomar et al., 2019; Singh et al., 2020). As such, a registration pipeline factoring in the internal organization of the brainstem would improve the validity of BOLD signal sampling at the individual level.

In principle, the quality of conventional standard-to-native, T1w-based registration could be enhanced by incorporating an additional image modality that provides spatial references to the internal architecture of the brainstem. Diffusion-weighted imaging (DWI) is one such modality that is well suited for this purpose. Unlike susceptibility-weighted imaging or turbo spin echo imaging, which are primarily used to visualize iron-rich structures localized to the midbrain or neuromelanin-laden nuclei distributed in a patchy manner, DWI-derived images can reveal the densely interwoven axonal tracts that span nearly the entire brainstem. Aside from its well-established role in identifying anatomical substrates for functional connectivity (van den Heuvel & Sporns, 2013), the microstructural features of axonal tracts revealed by DWI also form an internal spatial reference frame that could be used to improve ROI registration accuracy for the brainstem nuclei embedded within it. Typically, DWI enables the derivation of various scalar and vector maps—such as the fractional anisotropy (FA), the non-diffusion-weighted (b0), the mean diffusivity (MD), and the fiber direction maps—which offer complementary microstructural information relevant for ROI delineation. These DWI-derived maps are inherently co-registered to each other, thereby avoiding registration errors that can arise when aligning images obtained from different MR sequences.

In this study, we evaluated the benefit of incorporating DWI-derived images in addition to the T1w for brainstem nuclei ROI registration. Specifically, standard-to-native transforms were generated through a two-step process that first computed a whole-brain T1w-based transform, as in the conventional approach, and then locally refined the transform within the brainstem by co-registering (i) the T1w image with a T1w template, (ii) the FA image with an FA template, (iii) the b0 image with a b0 template, (iv) all three image-template pairs in (i)-(iii) through an iterative multi-channel approach. These transforms were subsequently used to map brainstem ROIs from standard space into the native spaces of individual participants. The performance of the four types of transforms—T1w-based, FA-based, b0-based, and multivariate—was evaluated by quantifying the spatial overlap between the registered ROIs and manually delineated brainstem ROIs, which served as the ground truth. We hypothesized that incorporating DWI-derived images, by introducing internal structural features of the brainstem, would improve the accuracy of brainstem ROI registration over conventional T1w-based registration.

## 2 Materials and methods

### 2.1 Participants

Data were acquired from twenty healthy participants (age: 42.15 ± 15.71 years; ten females) using a Siemens Magnetom Prisma 3 T scanner (Siemens Healthineers, Erlangen, Germany) at Toronto Western Hospital, equipped with a community-standard 20-channel head and neck receive coil. The study was approved by the University Health Network Research Ethics Board (REB 21-6105), and signed informed consent was obtained from all participants.

### 2.2 Data acquisition

T1-weighted (T1w) images were acquired using a 3D magnetization-prepared rapid acquisition gradient echo (MPRAGE) sequence with the following parameters (Mugler & Brookeman, 1990): matrix size = 320 × 320 × 160; resolution = 0.65 × 0.65 × 1 mm^3^; echo time (TE) = 2.6 ms; repetition time (TR) = 2200 ms; inversion time (TI) = 944 ms; flip angle = 8°; GRAPPA = 2; bandwidth = 252 Hz/pixel. Diffusion-weighted images (DWI) were acquired using a spin-echo SMS 2D-EPI sequence in two anterior-posterior (AP) runs: one with 64 diffusion directions (b = 2000 s/mm²) and one with 30 directions (b = 1000 s/mm²). Each run included five b0 images. An additional five b0 images were acquired with posterior-anterior (PA) phase encoding for distortion and eddy corrections. Acquisition parameters were: GRAPPA=2, SMS=2, resolution = 2 × 2 × 2 mm³; matrix size = 110 × 110 × 76; TE = 75 ms; TR = 4100 ms; flip angle = 90°. Susceptibility-weighted images were acquired using a multi-echo gradient-echo sequence: matrix size = 384 × 384 × 52; resolution = 0.5 × 0.5 × 1 mm³; TR = 53 ms; four echoes (TE = 7.3, 20, 32, 44 ms); flip angle = 20°; bandwidth = 100 Hz/pixel. All the sequences and parameter set are available at: https://zenodo.org/records/10685449

### 2.3 Data preprocessing

DWI data were denoised using the MP-PCA algorithm implemented in the `dwidenoise` tool in MRtrix3. Ten AP b0 images and five PA b0 images were used to estimate susceptibility-induced off-resonance fields via FSL’s `topup` tool (Andersson et al., 2003). These outputs were passed to the `eddy` tool along with diffusion parameters to correct for eddy current-induced distortions, magnetic field inhomogeneities, and motion (Andersson et al., 2016) in a single-step. The corrected DWIs and rotated b-vectors were used with FSL’s `dtifit` to compute the diffusion tensor and scalar maps, including the fractional anisotropy (FA) and averaged b0 images. SWI data were processed using CLEAR-SWI, which optimizes multi-echo combination of phase and magnitude images to yield optimal susceptibility-based tissue contrast (Eckstein et al., 2021).

### 2.4 Image co-registration

The Brainstem Navigator Atlas v0.9 (https://www.nitrc.org/projects/brainstemnavig/) was selected for this study as it includes both structural and diffusion templates in the IIT space. All three IIT templates used in the study—namely, IIT-T1w, IIT-FA, and IIT-b0—were brain-extracted using a whole-brain mask derived from the IIT-T1w template to remove extra-brain structures that could interfere with registration (e.g., the bright basilar artery in IIT-T1w).

Within-participant alignment of the T1w, FA, and b0 images was performed in several steps. First, each participant’s T1w image was brain-extracted using FreeSurfer’s `mri_synthstrip` and then bias-field corrected using the `N4BiasFieldCorrection` tool in ANTs (Hoopes et al., 2022; Tustison et al., 2010; Tustison et al., 2021; Avants et al., 2009). The T1w, FA, and b0 images were resampled to a common resolution (0.7 × 0.7 × 0.7 mm³) to prevent resolution mismatches from affecting image alignment. Given that b0 images more closely resemble T1w images in terms of gross anatomical contrast (albeit inverted), the affine transform aligning diffusion and structural images was estimated between the b0 and T1w images using ANTs’ `antsRegistration` tool, with mutual information (MI) as the optimization metric. This transform was subsequently applied to the FA and b0 images to align them with the T1w anatomical space. To assess the robustness of our findings to potential interpolation artifacts, we repeated the entire analysis using images resampled to an isotropic resolution of 1 × 1 × 1 mm³. The results of this sensitivity analysis are provided in the Supplementary Material (Figure S2).

Brainstem ROIs were mapped from IIT space to each participant’s native space using compound transforms derived from a two-step registration pipeline illustrated in Figure 1A. In the first step, a T1w-based whole-brain standard-to-native transform (T_T1w-WB_) was estimated using ANTs’ `antsRegistration`, constrained to the participant’s whole-brain mask. This registration included sequential optimization of (1) a four-level rigid-body transformation using MI, (2) a four-level affine transformation using MI, and (3) a five-level symmetric normalization (SyN) transformation using cross-correlation (CC) as the metric (Avants et al., 2007). The T_T1w-WB_ transform was then applied to the IIT-T1w, IIT-FA, and IIT-b0 templates to create three warped templates in approximate alignment with the participant’s native space. Quality control was carried out using visual inspection for all stages of the registration process.

**Figure 1.**
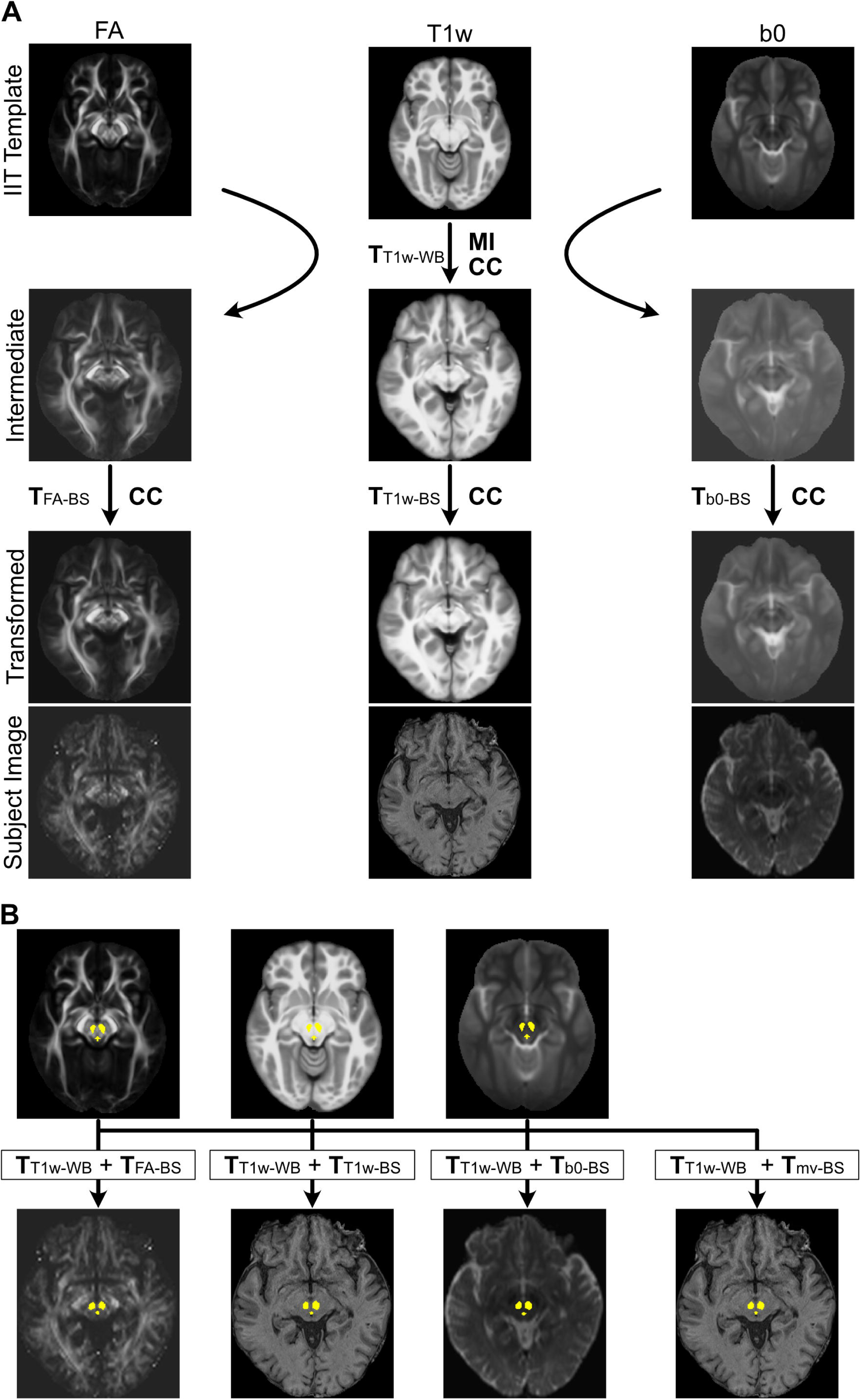
ROI registration procedure. (A) Estimation of transforms for the two-step registration pipelines. The whole-brain transform (T_T1w-WB_) was estimated via sequential linear (mutual information, MI) and nonlinear (cross-correlation, CC) alignment between the IIT-T1w template and the participant’s native T1w image. T_T1w-WB_ was then applied to the IIT-FA and IIT-b0 templates to generate warped intermediate templates. Brainstem-specific transforms (T_FA-BS_, T_T1w-BS_, and T_b0-BS_) were subsequently estimated by CC-based nonlinear registration between each warped template and its corresponding participant image. (B) ROI registration using compound transforms. The transforms obtained in (A) were concatenated to form four compound transforms—either including or excluding diffusion-based alignment—which were then applied to register brainstem ROIs from IIT space to native space.

In the second step, ANTs’ `antsRegistration` was used to estimate three additional nonlinear transforms by registering each warped IIT template (T1w, FA, b0) to its corresponding native image within the brainstem. A multivariate transform was also computed by jointly optimizing alignment across all three image types with equal weighting. All second-step registrations were constrained to a slightly dilated brainstem mask generated using FreeSurfer’s `recon-all` to ensure inclusion of relevant surrounding contours. The registrations were optimized using a CC-based, two-level SyN algorithm with the following parameters: convergence = 50 × 20; shrink factors = 2 × 1; smoothing sigmas = 2 × 0. The resulting nonlinear transforms—T_T1w-BS_, T_FA-BS_, T_b0-BS_, and the multivariate T_mv-BS_—were each concatenated with T_T1w-WB_ to form compound transforms for mapping brainstem ROIs from IIT space to each participant’s native space (Figure 1B).

The choice of ANTs for constructing the registration pipelines was motivated by prior benchmarking studies demonstrating its superior performance—specifically, the SyN algorithm—relative to other widely used non-linear registration methods, including FSL FNIRT, SPM DARTEL, and SPM Unified Segmentation (Klein et al., 2009; Avants et al., 2008; Avants et al., 2011). In addition, ANTs provides native support for multivariate registration, which constitutes a critical component of our approach. Registration scripts of this study can be accessed at: https://github.com/yianchenmil/BrainstemReg

### 2.5 Brainstem ROI delineation

Three sets of brainstem ROIs were selected to evaluate registration accuracy: the red nucleus (RN), substantia nigra (SN), and dorsal raphe nucleus (DRN). These ROIs were chosen based on the clear visibility of either their own anatomical boundaries or those of adjacent structures in native space, allowing for reliable comparison between registered ROIs and manually delineated ground-truth references. ROI delineation was performed using ITK-SNAP and 3D Slicer by YAC and YK. Both raters are medical doctors with formal training in neuroanatomy; YAC is a licensed general practitioner, and YK is a board-certified neuroradiologist.

The RN appears as a pair of low-intensity, ellipsoidal structures in the midbrain and was delineated based on its clearly identifiable boundaries in both susceptibility-weighted and T2-weighted b0 images. The SN is identified as a pair of low-intensity, crescent-shaped sheets located antero-inferolaterally to the RN in susceptibility-weighted and T2-weighted images; in particular, the boundary between the subthalamic nucleus and the SN is discernible in coronal susceptibility-weighted images. The well-circumscribed morphology of the RN and SN makes them suitable for manual delineation and Dice coefficient–based evaluation of registration accuracy.

In contrast, the DRN is not directly visualizable in any available MRI contrast and is therefore unsuitable for manual delineation or Dice coefficient–based assessment. However, the DRN lies directly ventral and parallel to the cerebral aqueduct and the fourth ventricle, both of which are clearly identifiable in T1-weighted images. Accordingly, we developed an alternative metric—the mis-registration fraction—to assess DRN registration accuracy. Instead of delineating the DRN itself, ROIs of the cerebral aqueduct and fourth ventricle (CA–4thVen) were delineated and used to compute this metric. Specifically, the mis-registration fraction quantifies the proportion of voxels within the registered DRN ROI that erroneously overlap with voxels of the cerebral aqueduct and fourth ventricle. Ground-truth ROIs for the cerebral aqueduct and fourth ventricle (CA–4thVen) were generated using FreeSurfer’s `recon-all` pipeline, with manual editing performed as needed to correct missing or incomplete segmentation of the narrow cerebral aqueduct.

It should be noted that although all three sets of brainstem ROIs were analyzed, only the results for the RN and DRN are presented in the main Results section, whereas the SN results are reported in the Supplementary Materials (Figure S1). This organization was adopted for two reasons. First, despite complete coverage of the SN in the b0 images, the caudal portion of the SN is not sufficiently captured in the susceptibility-weighted images for a few participants, raising concerns about methodological consistency across participants. Second, the SN ROI provided by Brainstem Navigator shows substantial deviation from the hypointense regions at the caudal and rostral ends in the IIT-b0 template (Figure S1A), raising concerns regarding the validity of this standard ROI. We therefore restrict the main presentation to results derived from ROIs that are better validated. Nevertheless, the SN results are included in the Supplementary Materials for interested readers.

### 2.6 Data analysis

The quality of brainstem ROI registration achieved by the four compound transforms (T1w-based, FA-based, b0-based, and multivariate) was evaluated using two metrics, as illustrated in Figure 2. First, the Dice coefficient between each registered RN ROI and the corresponding ground-truth ROI was calculated as:

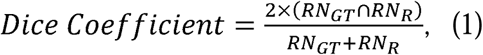

**Figure 2.**
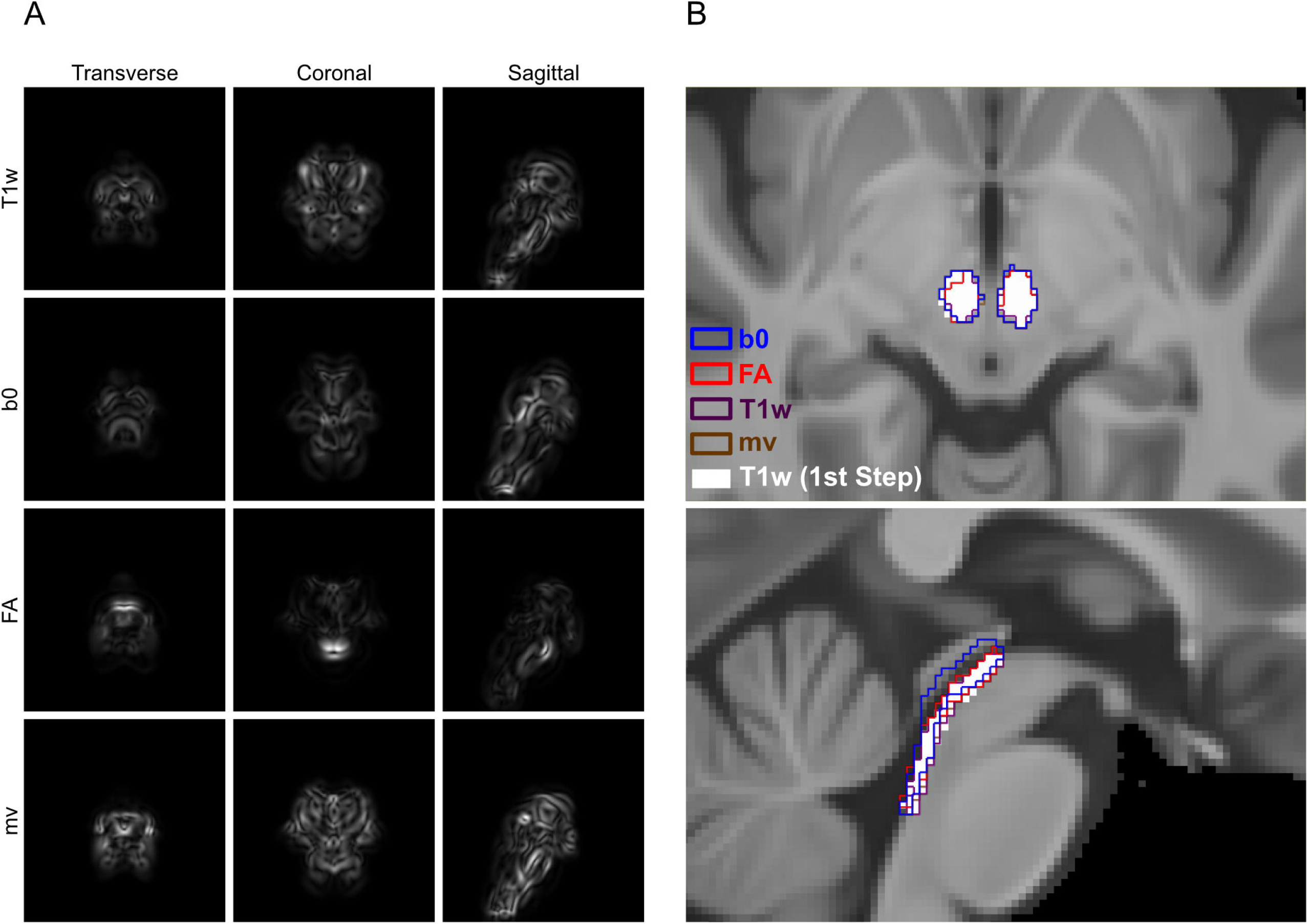
Evaluation metrics for assessing registration accuracy of the red nucleus (RN) and dorsal raphe nucleus (DRN). The Dice coefficient was used to evaluate RN registration accuracy by quantifying the spatial overlap between the registered ROI and the manually delineated ground-truth ROI. It is calculated as the ratio of twice the intersection volume to the sum of the two ROI volumes. For DRN, registration accuracy was assessed using the mis-registration fraction, defined as the proportion of voxels in the transformed DRN ROI that overlapped with the ground-truth ROI of the cerebral aqueduct and fourth ventricle.

where RN_GT_ denotes the ground-truth RN ROI and RN_R_ denotes the registered RN ROI. The Dice coefficients of SN ROIs were calculated in the same way.

Second, registration error for DRN was evaluated using the mis-registration fraction, defined as:

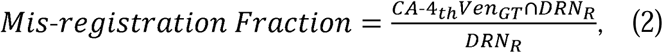

where DRN_R_ is the registered DRN ROI and CA-4_th_Ven_GT_ is the ground-truth CA-4_th_Ven ROI. For each participant, eight Dice coefficients—four for the RN and four for the SN—and four mis-registration fractions were determined, reflecting registration performance under the four different registration strategies across distinct brainstem regions. All analyses were conducted using Python 3 with `nibabel` for image processing.

### 2.7 Data visualization

To qualitatively compare the registration approaches, two types of visualizations were generated. First, four sets of probabilistic maps for RN, SN and DRN were created in IIT space by inverse-transforming the native-space ROIs obtained via the two-step compound transforms, using each participant’s inverse T_T1w-WB_ transform. These inverse-transformed ROIs were averaged across participants to generate probabilistic maps, which were compared with the original Brainstem Navigator atlas. It was hypothesized that diffusion-based second-step transforms (i.e., T_FA-BS_, T_b0-BS_, and the multivariate T_mv-BS_), by incorporating internal brainstem architecture, would result in greater spatial dispersion of the probabilistic maps due to the known inter-individual variability in brainstem ROI locations, as reported in previous studies with a similarly sized sample (Bianciardi et al., 2018; García-Gomar et al., 2019; Singh et al., 2020).

Second, Jacobian determinant maps (JDet) were computed for each second-step transform using ANTs’ `CreateJacobianDeterminantImage` tool. These maps were normalized to the range [0, 1], and gradient magnitude maps (|∇JDet|) were derived by calculating the spatial gradients and taking the magnitude of each voxel’s gradient vector. The resulting |∇JDet| maps were then averaged across participants to generate mean |∇JDet| maps, which were used to visualize regions of deformation targeted by the optimization algorithm. Since deformation tends to occur near modality-specific anatomical landmarks, such as the brainstem surface in T1w images and tract boundaries in FA images, these regions were expected to show higher gradient magnitudes in the corresponding |∇JDet| maps. The mean |∇JDet| map for the multivariate transform was expected to exhibit a more spatially uniform gradient distribution, reflecting the integration of salient anatomical features from all three image modalities.

## 3 Results

Figure 3A shows the Dice coefficients between the ground-truth RN ROIs and the four RN ROIs obtained via the two-step compound transforms for each of the twenty participants. A one-way repeated-measures ANOVA was conducted to examine the effect of compound transform type on Dice coefficient values. The analysis revealed a statistically significant difference among the four groups (F(3, 57) = 61.3106, p < 0.001). Post hoc pairwise t-tests with Bonferroni correction (adjusted α = 0.0083, 6 comparisons) indicated that Dice coefficients derived from the T1w-based compound transforms were significantly lower than those from the other three transform types (t(19) = -8.40, p < 0.001 for T1w versus b0; t(19) = -10.321, p < 0.001 for T1w versus FA; t(19) = -11.44, p < 0.001 for T1w versus multivariate), suggesting improved registration accuracy when diffusion-based images were incorporated into the registration process. In addition, the difference between the b0-and FA-based transforms was also statistically significant (t(19) = −3.31, p = 0.004). A parallel analysis of SN ROIs yielded similar results, indicating improved registration accuracy with the inclusion of diffusion-based images, particularly FA images. However, for the reasons outlined in Section 2.5 of the Materials and Methods, the SN results are presented in the Supplementary Material (Figure S1, Table S1).

**Figure 3.**
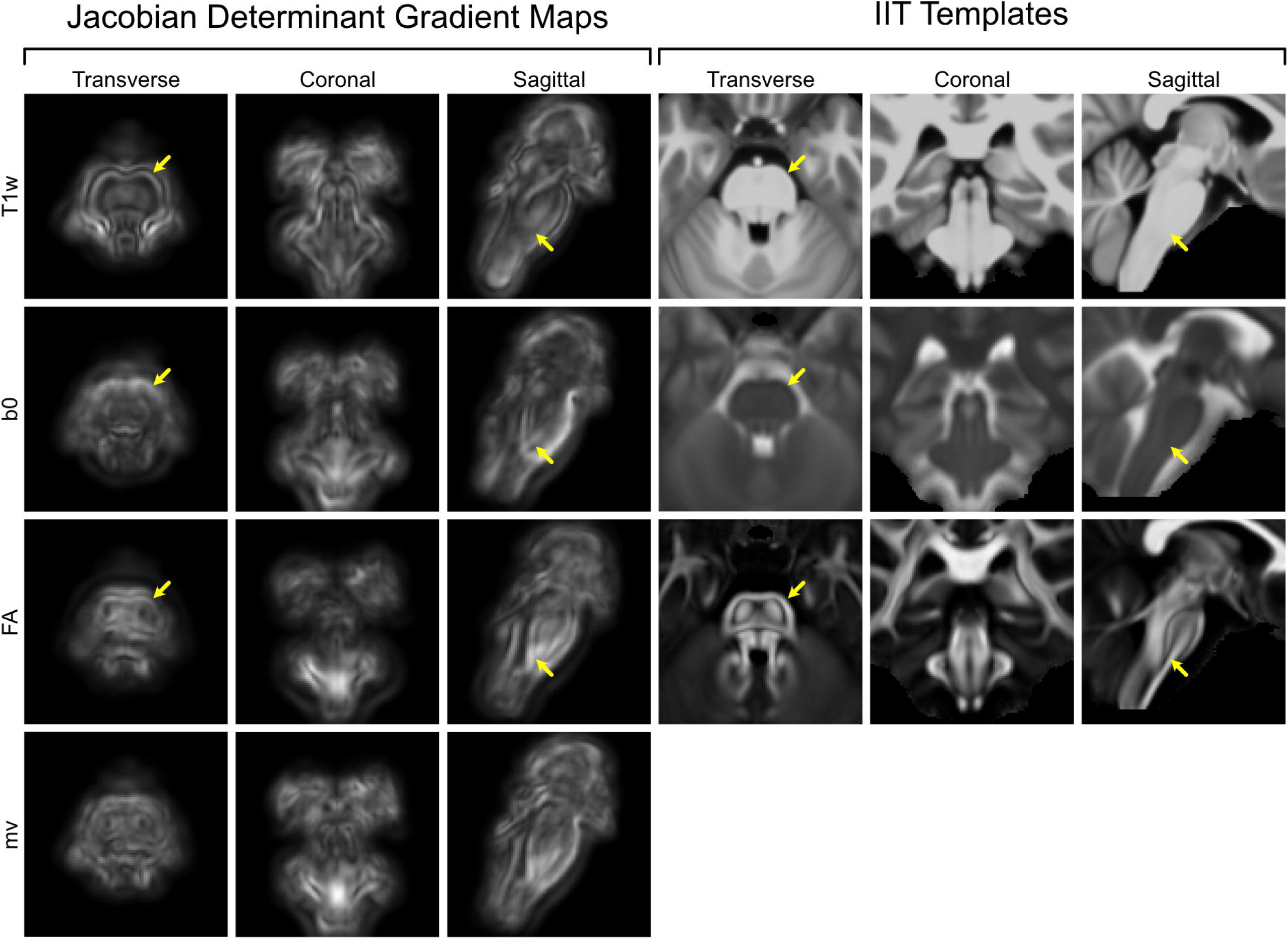
Registration accuracy of the four ROI registration pipelines. (A) Dice coefficients for the red nucleus (RN) across twenty participants, with each participant represented by a separate colored line. Mean ± standard deviation: T1w = 0.74 ± 0.02; b0 = 0.77 ± 0.02; FA = 0.79 ± 0.02; multivariate (mv) = 0.78 ± 0.02. (B) Mis-registration fractions for the dorsal raphe nucleus (DRN) across the same participants. Mean ± standard deviation: T1w = 0.21 ± 0.07; b0 = 0.23 ± 0.12; FA = 0.10 ± 0.06; mv = 0.16 ± 0.07.

Figure 3B shows the mis-registration fractions computed between the ground-truth CA-4_th_Ven ROIs and the four DRN ROIs obtained via the compound transforms for each of the twenty participants. A one-way repeated-measures ANOVA was conducted to examine the effect of compound transform type on mis-registration fractions. The analysis revealed a statistically significant difference among the four groups (F(3, 57) = 26.61, p < 0.001). Post hoc pairwise t-tests with Bonferroni correction were conducted for all pairwise comparisons. The FA-based compound transforms, which exhibited the lowest mean mis-registration fraction, differed significantly from each of the other three transform types (T1w vs. FA: t(19) = 8.82, p < 0.001; b0 vs. FA: t(19) = 6.29, p < 0.001; FA vs. multivariate: t(19) = −8.81, p < 0.001). The multivariate transform also significantly outperformed the T1w- and b0-based transforms, whereas no statistically significant difference was observed between the b0- and T1w-based transforms. All statistical results are summarized in Table 1. As a sensitivity analysis, the entire procedure was repeated using images resampled to 1 mm³ resolution; the corresponding results are presented in Figure S2 and Table S2.

**Table 1.** Summary statistics of registration accuracy metrics of RN and DRN.

| Comparison | Test | df | Statistic | p value | Significance |
| --- | --- | --- | --- | --- | --- |
| Four Pipelines | RM-ANOVA | (3, 57) | F=61.3106 | p<0.001 | Yes |
| T1w v.s. b0 | T test | 19 | T=-8.4036 | p<0.001 | Yes |
| T1w v.s. FA | T test | 19 | T=-10.3218 | p<0.001 | Yes |
| T1w v.s. mv | T test | 19 | T=-11.4355 | p<0.001 | Yes |
| b0 v.s. FA | T test | 19 | T=-3.3132 | p=0.0037 | Yes |
| b0 v.s. mv | T test | 19 | T=-2.8250 | p=0.0108 | No |
| FA v.s. mv | T test | 19 | T=2.3159 | p=0.0319 | No |

DRN
| Comparison | Test | df | Statistic | p value | Significance |
| --- | --- | --- | --- | --- | --- |
| Four Pipelines | RM-ANOVA | (3, 57) | F=26.6104 | p<0.001 | Yes |
| T1w v.s. b0 | T test | 19 | T=-0.8927 | p=0.3832 | No |
| T1w v.s. FA | T test | 19 | T=8.8210 | p<0.001 | Yes |
| T1w v.s. mv | T test | 19 | T=4.5533 | p=0.0002 | Yes |
| b0 v.s. FA | T test | 19 | T=6.2905 | p<0.001 | Yes |
| b0 v.s. mv | T test | 19 | T=3.6799 | p=0.002 | Yes |
| FA v.s. mv | T test | 19 | T=-8.8116 | p<0.001 | Yes |

Figure 4 presents probabilistic maps of the DRN (top row) and RN (bottom row), generated by inverse-transforming the two-step-transformed native-space ROIs of each participant back into IIT space and averaging across the twenty participants. For reference, the unthresholded Brainstem Navigator probabilistic maps for each nucleus are shown in the leftmost column. In every panel, red contours mark the thresholded Brainstem Navigator ROIs (threshold = 0.35), which served as the standard ROIs in IIT space throughout the registration pipelines.

**Figure 4.**
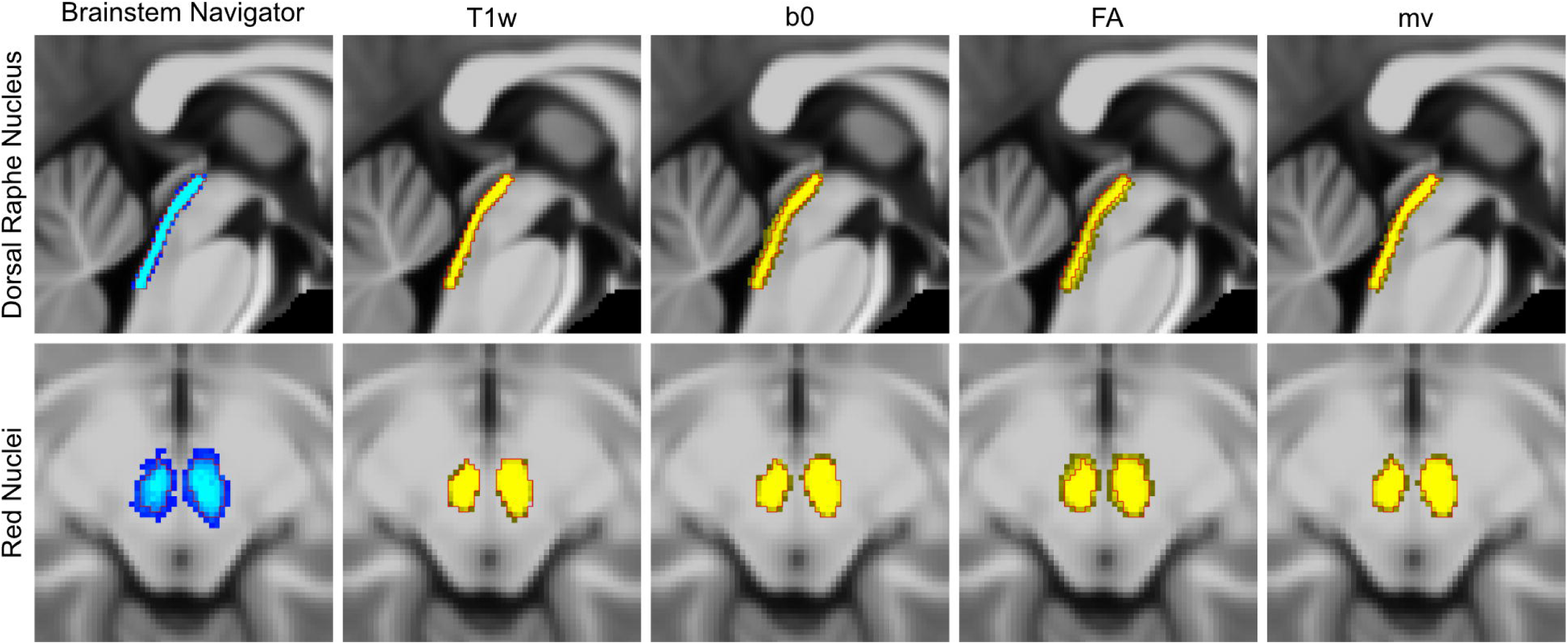
Probabilistic ROIs of the red nucleus (RN) and dorsal raphe nucleus (DRN). The leftmost column shows the unthresholded probabilistic ROIs from the Brainstem Navigator atlas (blue), overlaid with the boundaries of the thresholded binary ROIs at probability = 0.35 (red). Columns from second-to-left through rightmost display the probabilistic ROIs constructed by averaging inverse-transformed native-space ROIs obtained from each of the four two-step registration pipelines (yellow), overlaid with the same thresholded ROI boundaries (red).

As shown in the Brainstem Navigator maps, only 68.1% of voxels in the DRN probabilistic map and 51.2% in the RN map fall within the thresholded ROI boundaries. These relatively low proportions reflect the anatomical variability among individuals whose brainstem data were used to construct the original probabilistic atlas, as shown in the original studies (Bianciardi et al., 2018; García-Gomar et al., 2019; Singh et al., 2020). In contrast, the T1w-based probabilistic ROIs showed much higher containment: 89.9% of DRN and 87.1% of RN voxels fell within the thresholded boundaries. This suggests that augmenting the whole-brain T1w-based transform (T_T1w-WB_) with a local T1w-based refinement (T_T1w-BS_) contributes little additional individual specificity; the resulting two-step transformed ROIs remain largely consistent with the conventional one-step method.

By comparison, the DWI-augmented probabilistic ROIs show a broader spatial distribution, with smaller proportions of their voxels falling within the thresholded boundaries, namely 58.9% for b0-augmented DRN, 49.0% for FA-augmented DRN, and 65.8% for multivariate-augmented DRN; and 80.7%, 59.6%, and 73.9% for the corresponding RN maps. Visually, these ROIs appear less tightly clustered and more dispersed across subjects. This increased dispersion suggests that the DWI-augmented registration approaches better capture individual anatomical variability and produce more personalized ROI mappings. A parallel analysis of SN ROIs yielded similar voxel proportion distributions, with the results shown in Figure S1C.

Figure 5 presents the mean gradient magnitude maps of the Jacobian determinants (|∇JDet|) derived from the four second-step transforms, displayed in transverse, coronal, and sagittal planes. Corresponding IIT templates (T1w, b0, and FA) are shown on the right for anatomical reference. As indicated by the yellow arrows in the transverse plane, the T1w-based |∇JDet| map exhibits a high-intensity rim that closely matches the outer contour of the brainstem visible in the IIT-T1w template. This pattern suggests that alignment during the T1w-based registration process was primarily driven by the brainstem surface. The interior regions of the T1w-based |∇JDet| map display relatively low gradient magnitudes, though a faint symmetrical pattern implies some internal alignment.

**Figure 5.**
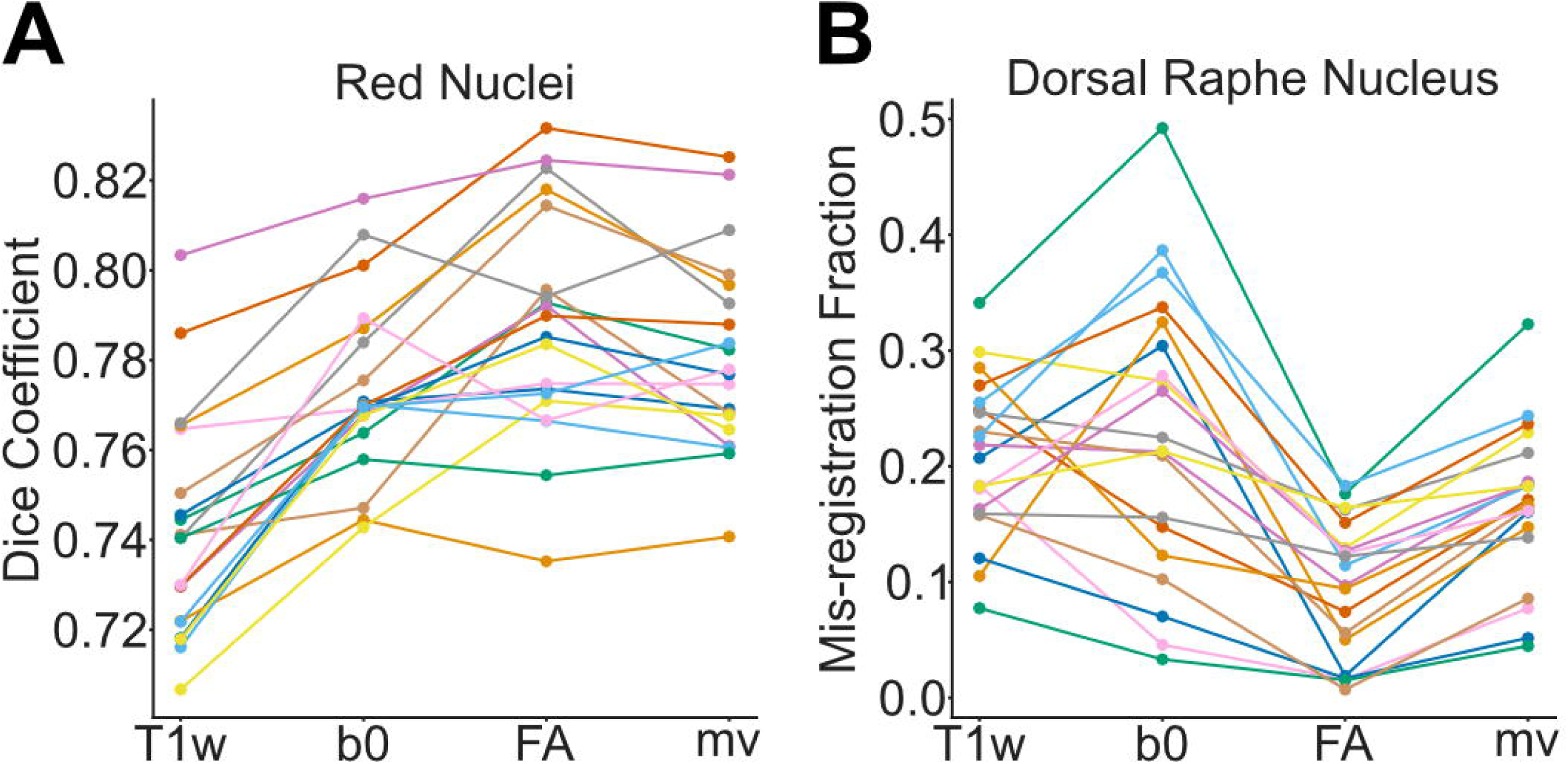
Gradient magnitude maps of brainstem-specific transforms and corresponding templates. Left: Mean gradient magnitude maps of the Jacobian determinants (|∇JDet|) computed from each brainstem-specific transform, shown in transverse, coronal, and sagittal planes. Right: Corresponding IIT templates (T1w, b0, and FA) displayed in the same anatomical planes for reference. Yellow arrows highlight subregional differences in deformation magnitude, which may reflect variation in anatomical landmark visibility across image modalities.

In contrast, the b0- and FA-based |∇JDet| maps show less emphasis on the outer boundary, as marked in the corresponding locations. Instead, both maps exhibit multiple symmetric, high-magnitude clusters within the brainstem, indicating enhanced sensitivity to internal anatomical features. This is particularly evident at the pontomedullary junction in the sagittal plane, where the T1w-based map remains dark while the FA-based map shows localized bright gradients. This contrast likely reflects underlying template differences: the IIT-T1w template is relatively homogeneous in this region, whereas the IIT-FA template reveals intersecting fiber tracts at the pontomedullary junction.

The multivariate |∇JDet| map combines features observed in the unimodal maps, showing gradient magnitudes that are more uniformly distributed across the brainstem. This pattern suggests that the multivariate transform leveraged both external and internal anatomical features from the three image modalities.

## 4 Discussion

In this study, we evaluated several DWI-augmented ROI registration pipelines for their effectiveness in mapping standard brainstem ROIs from IIT space to individual participants’ native spaces, with the design of the pipelines and associated evaluation metrics summarized in Figures 1 and 2. As shown in Figures 3 and S1, DWI-derived images improved registration accuracy in a manner that varied by both ROI and image modality. The greater spatial dispersion observed in the probabilistic maps of inverse-transformed ROIs, as shown in Figure 4, suggests that DWI-augmented pipelines may better capture inter-individual anatomical variability. The |∇JDet| maps in Figure 5 further reveal how different pipelines distribute registration effort across brainstem subregions, offering a qualitative explanation for the observed ROI- and modality-dependent effects. These findings have important implications for the brainstem fMRI community, which continues to face technical challenges, particularly the unreliable sampling of brainstem signals resulting from inaccurate localization of target ROIs. The findings could also benefit research areas that rely heavily on accurate data sampling from small brainstem ROIs, such as brainstem histopathology.

There are, however, some limitations and issues in this study that would warrant further investigation. One key issue concerns the interpretation of the increased spatial dispersion observed in the DWI-augmented probabilistic ROIs shown in Figure 4. This dispersion may reflect enhanced sensitivity of the DWI-augmented pipelines to interindividual anatomical variability. Alternatively, it could arise from reduced registration accuracy attributable to the lower signal-to-noise ratio and greater geometric distortion of diffusion images. The overall pattern of data presented in Figure 3 and S1 suggests that individual anatomical variability is the more likely explanation for two reasons. First, the DWI-augmented pipelines appear to modulate within-group variability of the accuracy metrics across all three sets of brainstem ROIs in a structure- and modality-dependent manner. For example, the b0-based pipeline decreases within-group variability for the RN while increasing it for the DRN, whereas DRN within-group variability increases with b0-based pipeline but decreases with FA-base pipeline. This pattern is more consistent with DWI-based pipelines achieving individualized, structure- and modality-specific alignments than with a general failure of registration attributable to diffusion image quality. Second, the Dice coefficients for the RN and SN shown in Figure 3 and Figure S1 indicate improved, rather than diminished, accuracy with the DWI-augmented pipelines. This finding suggests that voxels are aligned to their correct, individualized spatial locations rather than being dispersed randomly. For DRN registration, however, the possibility that diffusion images introduce additional registration uncertainty cannot be entirely excluded. The mis-registration fraction metric captures voxel displacement only in the posterior direction relative to the DRN and does not account for misalignment in other directions. Consequently, the reduced mis-registration fraction achieved by FA-based and multivariate transforms reflects improved registration accuracy limited to the dorsal aspect of the DRN. Accordingly, we interpret the observed spatial dispersion conservatively: it most likely reflects genuine anatomical variability rather than random error, although more comprehensive registration metrics across all evaluated nuclei will be required to draw a definitive conclusion, particularly given the potential for regional variation in diffusion image quality.

This lingering uncertainty stems from a broader limitation: the unavailability of MR sequences capable of reliably delineating the full boundaries of many brainstem nuclei at the individual level. As a result, alternative evaluation metrics that do not require complete ground-truth segmentation, such as the mis-registration fraction, must be used to assess registration accuracy for these structures. Although the Dice coefficient results support the interpretation that DWI-augmented pipelines yield more anatomically specific registrations, it would be preferable to validate this claim using nuclei that can be directly visualized and manually segmented in native space, once advanced MR sequences and ultra-high-field MRI scanners become more widely available. It is worth noting that even as such sequences are developed, the optimization of atlas-to-subject ROI registration pipelines will remain important, as incorporating multiple specialized scans to visualize different brainstem nuclei may not always be practical in research or clinical settings due to limitations in scan time, participant motion, and subject tolerance (Greene et al., 2016).

Another issue warranting further systematic exploration—though beyond the scope of the present work—is the refinement of the multivariate registration pipeline. As described in the Materials and Methods section, the T_mv-BS_ transforms were estimated using equally weighted image modalities, which is the default setting in ANTs’ `antsRegistration` tool for multivariate optimization. However, the optimal weighting may vary across subregions or specific brainstem nuclei, given differences in how well individual structures and their surrounding landmarks are visualized in each modality. For example, Figure 3 shows that incorporating b0-based optimization consistently improves the registration accuracy of the RN but appears to introduce greater variability in the mis-registration fraction of the DRN. This discrepancy likely stems from how well each structure can be visualized: while the b0 image provides direct contrast for the RN, it offers poor visibility in the dorsal midbrain, where the DRN is located. In contrast, the impact of FA-based optimization on DRN registration appears more consistent across participants, likely due to the presence of nearby salient white matter tracts in FA images. These observations suggest that tailoring modality weights based on prior anatomical knowledge could improve registration performance in an ROI-specific manner. Further investigation in this direction may enhance the flexibility and accuracy of the proposed multivariate pipeline across different brainstem subregions.

The significance of our findings is best appreciated in the context of the development and application of the Brainstem Navigator-the standard-space atlas used to evaluate the DWI-augmented registration pipelines in this study. As described in a series of publications (Bianciardi et al., 2015; Bianciardi et al., 2018; García-Gomar et al., 2019; Singh et al., 2020; Singh et al., 2021), the development of the Brainstem Navigator was guided by the same theoretical assumption as the present work: that DWI-derived images provide detailed microstructural information suitable for delineating brainstem nuclei. The Brainstem Navigator team segmented these nuclei using 7-Tesla b0 and FA images from twelve participants via a semi-automated method combining k-means clustering and manual labeling. These DWI-derived images and segmented nuclei ROIs were then registered to the IIT diffusion imaging templates to construct a probabilistic atlas in IIT space. Additionally, the regions of interest (ROIs) were transformed into MNI152 space using a T1w-based IIT-to-MNI152 registration, with the IIT version intended for the diffusion imaging community and the MNI152 version for the fMRI community (Bianciardi et al., 2015). Accordingly, we observed a divergence in ROI registration strategies across studies utilizing the Brainstem Navigator. While DWI-based studies tend to use the IIT-space version for diffusion image-based ROI registration (García-Gomar et al., 2021), fMRI studies typically rely on the T1w-based ROI registration pipelines included in standard fMRI preprocessing toolkits, applying them to the MNI152-space version of the atlas (Zhang et al., 2025; Hansen et al., 2024).

Although the present study focuses on the application rather than the development of a brainstem atlas, its methodology partially overlaps with that of the Brainstem Navigator, allowing for informative comparisons. First, the Brainstem Navigator team delineated a greater number of brainstem nuclei using only b0 and FA images. Second, although not formally evaluated, their native-to-IIT registration appeared satisfactory using FA images alone (Bianciardi et al., 2015). These differences may be attributable to the higher image quality achieved with a 7-Tesla scanner compared to the 3-Tesla scanner used in the present study. Due to the lower spatial resolution and signal-to-noise ratio in 3-Tesla DWI data, we were only able to manually delineate the boundaries of the RN and SN ROIs with sufficient confidence, and we needed to employ a metric that did not require full boundary delineation to assess the registration accuracy of DRN ROI. Additionally, direct registration of our 3-Tesla DWI-derived images to the IIT diffusion templates yielded suboptimal results. Therefore, we adopted a two-step registration approach: initial global alignment was performed using T1w-based whole-brain images (T_T1w-WB_), followed by local refinement using DWI-based images through cross-correlation optimization (T_FA-BS_, T_b0-BS_, and T_mv-BS_).

An intriguing aspect of the Brainstem Navigator’s approach is that, although the developers acknowledged T1w images lacked sufficient contrast for delineating brainstem nuclei—and therefore excluded them during ROI segmentation—they nonetheless employed a T1w-based transformation to map the atlas from IIT space to MNI152 space (Bianciardi et al., 2015; Bianciardi et al., 2018; García-Gomar et al., 2019). While this decision was not explicitly justified, it likely reflects the absence, at the time, of a widely accepted diffusion imaging template in MNI152 space—the default normalization space used in popular fMRI preprocessing toolkits like SPM and FSL (Zhang & Arfanakis, 2018; Ashburner et al., 2014; Jenkinson et al., 2012). However, registration in neuroimaging is an inherently ill-posed problem, where the resulting transformation depends strongly on the chosen image modality and the cost function used during optimization (Lange et al., 2024). Because T1w images lack adequate contrast for brainstem nuclei, performing native-to-standard or standard-to-standard (e.g., IIT-to-MNI152) ROI registration using only T1w images may compromise the anatomical precision afforded by an atlas developed through DWI-based delineation. This concern is supported by our findings that DWI-augmented pipelines outperformed the T1w-only pipeline in the context of IIT-to-native registration. To extend this evaluation to standard-to-standard registration, we generated IIT-to-OMM mappings by registering the IIT-T1w, IIT-b0, and IIT-FA templates to the recently released OMM-1 (Oxford-MultiModal-1) T1w, T2w-FLAIR, and FA templates (Arthofer et al., 2024). These mappings were computed using the same pipelines applied to IIT-to-native registration in the current study, with the sole modification that the CC cost function for the T_b0-BS_ pipeline was replaced by MI to accommodate the contrast differences between b0 and T2w-FLAIR images. As shown in Figure 6A, the |∇JDet| maps of the four brainstem-masked transforms varied substantially, despite all being IIT-to-OMM mappings. Consistent with this, the transformed RN and DRN ROIs also showed spatial discrepancies across the different registration pipelines (Figure 6B), mirroring patterns observed in our IIT-to-native results. These differences underscore the impact of image modality on registration accuracy—a factor that appears to have been underappreciated in the development of brainstem atlases and brainstem fMRI preprocessing toolkits. Although preliminary, our findings suggest that incorporating DWI-derived images at every stage of the registration process could mitigate the loss of accuracy associated with contrast limitations in certain modalities. For example, normalizing participant data to IIT space—an approach commonly used by DWI researchers employing the Brainstem Navigator—avoids the intermediate T1w-only IIT-to-MNI152 registration and may improve registration accuracy of brainstem ROIs. This strategy would require additional diffusion acquisitions in an fMRI session; however, with appropriate EPI distortion matching, these data may also be utilized to enhance functional-to-structural co-registration, as previously proposed and tested with T1w images in selected studies (Renvall et al., 2016; Zhang et al., 2025).

**Figure 6.**
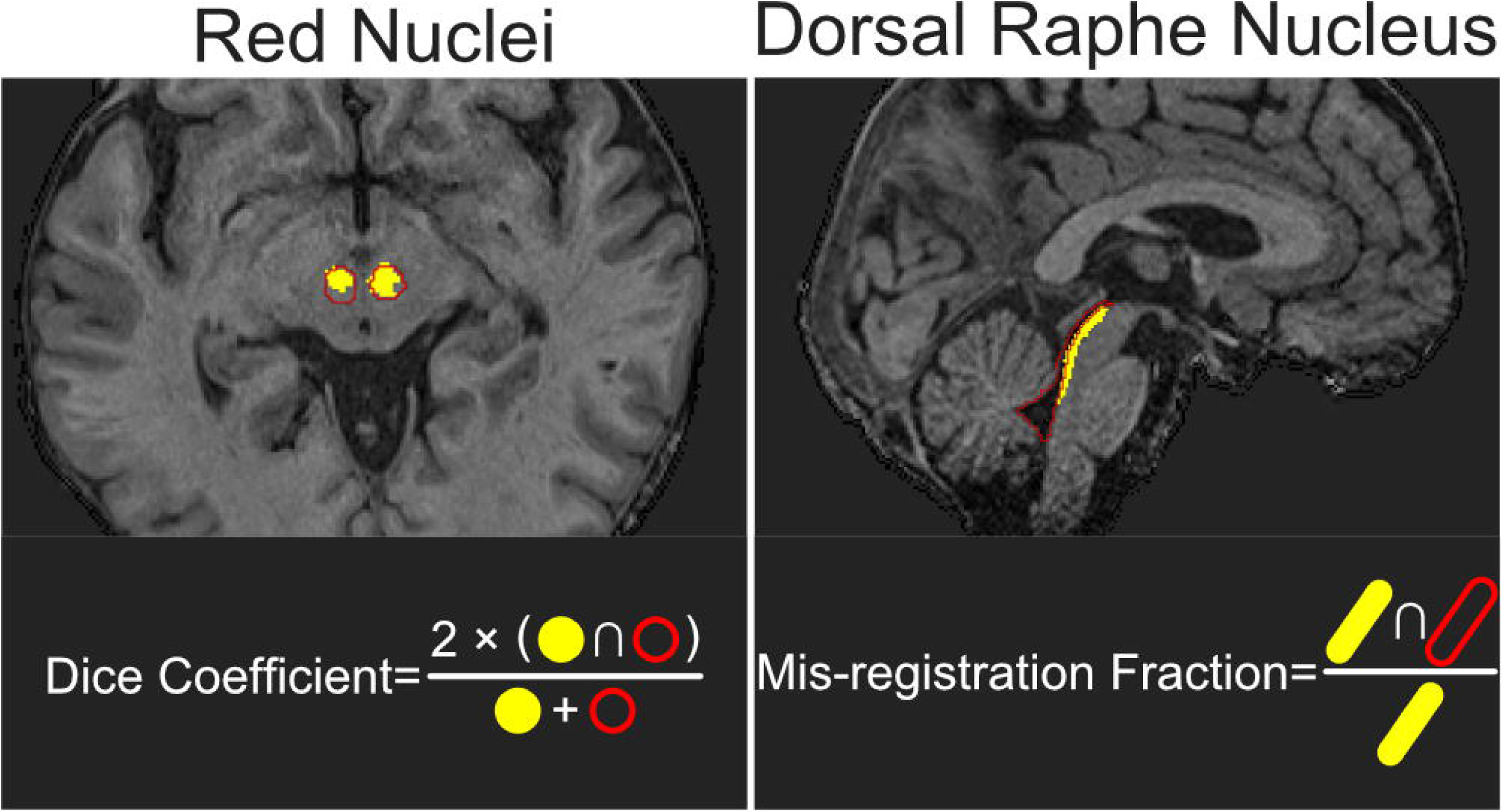
Gradient magnitude maps of brainstem-specific IIT-to-OMM transforms. (A) Gradient magnitude maps of the Jacobian determinants (|∇JDet|) computed from each brainstem-specific transform, shown in transverse, coronal, and sagittal planes. (B) Brainstem ROIs transformed from the IIT space to the OMM-1 space, overlaid on the OMM-1 T1w template. Upper: RN. Lower: DRN. b0: concatenated T_T1w-WB_ and T_b0-BS_. FA: concatenated T_T1w-WB_ and T_FA-BS_. T1w: concatenated T_T1w-WB_ and T_T1w-BS_. mv: concatenated T_T1w-WB_ and T_mv-BS_. T1w (1^st^ step): T_T1w-WB_ only.

An important question regarding the incorporation of DWI is how best to integrate it to maximize its benefits while minimizing potential deleterious effects, if such effects exist. The Brainstem Navigator atlas was constructed using a sequential approach, in which only one imaging modality was used at a time. For instance, the DRN was identified by first localizing the periaqueductal gray (PAG)-DRN complex on the b0 image, followed by delineation of DRN boundaries using the FA image. Similarly, b0 and FA images were used sequentially to delineate the outer boundaries and inner subregions of the RN, respectively (Bianciardi et al., 2015). The choice of modality for delineating specific structures was guided by prior knowledge of which image type provided superior contrast for each brainstem nucleus. Interestingly, our findings suggest that b0-based and FA-based registration produced more consistent improvements in registration accuracy for the RN and DRN respectively (Figure 3), paralleling the Brainstem Navigator team’s modality selection for ROI delineation. This suggests that the registration algorithms were able to leverage nucleus-specific structural information embedded in the corresponding DWI images to optimize the transformations. The Brainstem Navigator team also discussed the possibility of adopting a multi-channel approach, as opposed to a sequential one, for brainstem nuclei segmentation (Bianciardi et al., 2015). Although our study focused on registration rather than segmentation, informative comparisons can still be drawn from our results involving single-step (T_T1w-WB_), sequential (e.g., T_b0-BS_ and T_FA-BS_ following T_T1w-WB_), and multi-channel (T_mv-BS_ following T_T1w-WB_) registration pipelines, as illustrated in Figure 6B. Notably, the second-step T_b0-BS_ transform actually worsened DRN registration: more DRN ROI voxels were misplaced into the cerebral aqueduct and fourth ventricle compared to those registered using T_T1w-WB_ alone. This degradation was not observed with the multi-channel pipeline, in which FA, b0, and T1w images were simultaneously incorporated and iterated over within a single registration step. The deleterious effect observed with the T_b0-BS_ pipeline could be partly attributed to the use of MI rather than CC as the optimization metric due to subtle contrast mismatch between IIT-b0 and OMM-1 T2w-FLAIR. However, the fact that the degradation occurred specifically for the DRN suggests that it may also stem from the relatively poor contrast of these T2w images at the dorsal midbrain, where the DRN is located. This interpretation is consistent with the results shown in Figure 3B, which demonstrate lower Dice coefficients and increased within-group variability for b0-based DRN registration. In this case, the multi-channel registration approach, using default equal weighting across modalities, appears to effectively mitigate the adverse impact of the lower-contrast modality. As discussed earlier, optimizing the weighting of each modality may further enhance registration performance, but doing so would require prior knowledge of nucleus-specific contrast characteristics and systematic empirical evaluation. Importantly, once such optimized weighting schemes are established for brainstem ROI registration, the same principles could be translated to brainstem ROI segmentation. Since segmentation also relies on detecting subtle contrast differences between adjacent nuclei, tailoring the contribution of each modality based on its ability to resolve specific anatomical boundaries may improve the precision of segmentation algorithms, particularly in complex or low-contrast regions of the brainstem.

It should be noted that the DWI-augmented registration approach evaluated in this study was originally conceptualized for research contexts in which multiple brainstem nuclei need to be examined simultaneously (e.g., intra-brainstem functional connectivity analyses, as in Hansen et al., 2024). When the research focus is limited to a single brainstem structure that can be directly visualized using specialized MR sequences, we recommend prioritizing native visualization with those dedicated contrasts to achieve more precise localization. If DWI-augmented pipelines are applied to clinical research, the construction of study-specific templates and a multi-stage registration strategy incorporating group-specific templates (e.g., a Parkinson’s disease–specific template) would be necessary to mitigate biased registration accuracy arising from systematic structural differences between groups (Borghammer et al., 2010; Rao et al., 2017; Yang et al., 2020; Fillmore et al., 2015; Jia et al., 2011; Xiao et al., 2015). As noted above, specialized sequences should be prioritized for visualizing structures affected by pathological change, particularly when assumptions of topological equivalence between clinical and healthy groups may be violated (e.g., altered signal intensity due to iron deposition; see Pfefferbaum et al., 2010; Langley et al., 2021).

In summary, we proposed and evaluated a theoretically grounded approach to improving the registration accuracy of brainstem nuclei ROIs, addressing the persistent challenge posed by the limited availability of MRI sequences that enable direct visualization of these structures. Through a detailed methodological comparison with studies using Brainstem Navigator, we also offered cautious recommendations on how the anatomical precision provided by a brainstem atlas could be better preserved using a DWI-augmented registration pipeline tailored to specific brainstem regions. Despite certain limitations, most notably, the scarcity of ground-truth nuclei for rigorous validation, the results demonstrate both the feasibility of the approach and its potential for ROI-specific optimization. Although this study focused on two scalar images derived from DWI (b0 and FA), other diffusion-based images, such as the diffusion tensor image (DTI), contain rich, high-dimensional information that could support more precise local alignment and further augmentation with global anatomical features, such as tract labels from cortical-subcortical tractography. In the context of fMRI, cortical-subcortical tractography also provides a structural basis for investigating the mechanisms underlying functional dynamics aside from serving as a spatial reference frame for ROI registration, underscoring the value of incorporating DWI into fMRI protocols for enhancing fidelity and biological plausibility. As ultra-high field MRI becomes more widely available and methods for correcting geometric distortion continue to advance, the integration of diffusion-weighted data into ROI registration pipelines is likely to become increasingly sophisticated. Future research could explore these technical extensions and incorporate functional-to-structural co-registration for a more holistic enhancement of brainstem fMRI reliability.

## Acknowledgements

The authors thank Asma Naheed, MRT for the scanning support at the Slaight Family Centre for Advanced MRI, Toronto Western Hospital, University Health Network, Toronto, Canada. They also thank Dr. Chun-Yu Chen for his valuable advice on brainstem ROI delineation and for his guidance in addressing the clinical aspects of the reviewers’ comments at the Division of General Neurology, Neurological Institute, Taipei Veterans General Hospital, Taipei, Taiwan.

## Author Contributions

YAC: Conceptualization, Data curation, Formal analysis, Investigation, Methodology, Software, Visualization, Writing-original draft.

LK: Data curation, Writing-review & editing.

CTC: Data curation, Writing-review & editing.

YK: Data curation, Writing-review & editing.

AB: Data curation, Writing-review & editing.

JG: Data curation, Writing-review & editing.

AML: Funding acquisition, Resources, Writing-review & editing.

KU: Funding acquisition, Project administration, Resources, Supervision, Writing-review & editing.

AOD: Funding acquisition, Supervision, Writing-review & editing.

SK: Conceptualization, Data curation, Project administration, Resources, Software, Supervision, Writing-review & editing.

