## Supplementary Material for "Evaluating the quality of brainstem ROI registration using structural and diffusion MRI"

Figure S1. Registration accuracy of the four ROI registration pipelines and the corresponding probabilistic ROIs for the substantia nigra (SN). (A) Deviation of the thresholded binary Brainstem Navigator SN ROI (probability=0.35) from the hypointense regions in the IIT-b0 atlas. (B) Dice coefficients across twenty participants, with each participant represented by a separate colored line. Mean ± standard deviation: T1w = 0.46 ± 0.07; b0 = 0.50 ± 0.06; FA = 0.53 ± 0.07; multivariate (mv) = 0.50 ± 0.07. (C) Probabilistic ROIs of SN generated by averaging inverse-transformed native-space ROIs obtained from each of the four two-step registration pipelines (yellow). The unthresholded probabilistic ROIs from the Brainstem Navigator atlas is shown in the leftmost panel for comparison (blue). The red contour in each panel represents the boundary of the thresholded binary Brainstem Navigator ROI. Proportions of registered voxels within the boundary: 42.1% for the original unthresholded Brainstem Navigator probabilistic ROI, 70.9% for T1w, 66.9% for b0, 56.8% for FA, 64.0% for mv.

Figure S2. Registration accuracy of the four ROI registration pipelines created with images of 1 mm^3^ resolution. (A) Dice coefficients for the red nucleus (RN) across twenty participants, with each participant represented by a separate colored line. Mean ± standard deviation: T1w = 0.76 ± 0.03; b0 = 0.77 ± 0.02; FA = 0.78 ± 0.02; multivariate (mv) = 0.78 ± 0.02. (B) Mis-registration fractions for the dorsal raphe nucleus (DRN) across the same participants. Mean ± standard deviation: T1w = 0.23 ± 0.08; b0 = 0.23 ± 0.15; FA = 0.10 ± 0.07; mv = 0.18 ± 0.08.

Figure S3. Exploration of the effect of incorporating b0–T1w boundary-based registration (BBR) into the b0-based registration pipeline on dorsal raphe nucleus (DRN) registration accuracy. *b0* denotes the original b0-based pipeline. *BBR* denotes the modified pipeline in which a six-degree-of-freedom boundary-based registration—implemented using FSL FLIRT—between the warped IIT-b0 template and the native T1w image was used to initialize the second-step, brainstem-masked registration, with the aim of improving alignment of the cerebral aqueduct–fourth ventricle (CA–4thVen) ROI. This modification resulted in a mild but statistically significant improvement in registration accuracy (b0: 0.23 ± 0.12; BBR: 0.20 ± 0.11; *t*(19) = 8.12, *p* < 0.001).

Table S1. Summary statistics of registration accuracy metrics of SN

| Comparison | Test | df | Statistic | p value | Significance |
| --- | --- | --- | --- | --- | --- |
| Four Pipelines | RM-ANOVA | (3, 57) | F=37.8412 | p<0.001 | Yes |
| T1w v.s. b0 | T test | 19 | T=-5.1522 | p<0.001 | Yes |
| T1w v.s. FA | T test | 19 | T=-7.8450 | p<0.001 | Yes |
| T1w v.s. mv | T test | 19 | T=-6.5860 | p<0.001 | Yes |
| b0 v.s. FA | T test | 19 | T=-4.5100 | p=-0.0002 | Yes |
| b0 v.s. mv | T test | 19 | T=-1.8006 | p=0.0877 | No |
| FA v.s. mv | T test | 19 | T=6.6391 | p<0.001 | Yes |

Table S2. Summary statistics of registration accuracy metrics of RN and DRN, resolution = 1 mm^3^

RN

| Comparison | Test | df | Statistic | p value | Significance |
| --- | --- | --- | --- | --- | --- |
| Four Pipelines | RM-ANOVA | (3, 57) | F=16.5219 | p<0.001 | Yes |
| T1w v.s. b0 | T test | 19 | T=-1.9193 | p=0.0701 | No |
| T1w v.s. FA | T test | 19 | T=-5.4796 | p<0.001 | Yes |
| T1w v.s. mv | T test | 19 | T=-4.9137 | p<0.001 | Yes |
| b0 v.s. FA | T test | 19 | T=-3.7669 | p=0.0013 | Yes |
| b0 v.s. mv | T test | 19 | T=-5.0939 | p<0.001 | Yes |
| FA v.s. mv | T test | 19 | T=1.3233 | p=0.2014 | No |

DRN

| Comparison | Test | df | Statistic | p value | Significance |
| --- | --- | --- | --- | --- | --- |
| Four Pipelines | RM-ANOVA | (3, 57) | F=19.0300 | p<0.001 | Yes |
| T1w v.s. b0 | T test | 19 | T=-0.2874 | p=0.7769 | No |
| T1w v.s. FA | T test | 19 | T=10.1300 | p<0.001 | Yes |
| T1w v.s. mv | T test | 19 | T=6.4603 | p<0.001 | Yes |
| b0 v.s. FA | T test | 19 | T=4.5892 | p=0.0002 | Yes |
| b0 v.s. mv | T test | 19 | T=2.5164 | p=0.02 | No |
| FA v.s. mv | T test | 19 | T=-8.1659 | p<0.001 | Yes |
