## Supplementary figures and images for "Evaluating the quality of brainstem ROI registration using structural and diffusion MRI"

### Figure S1

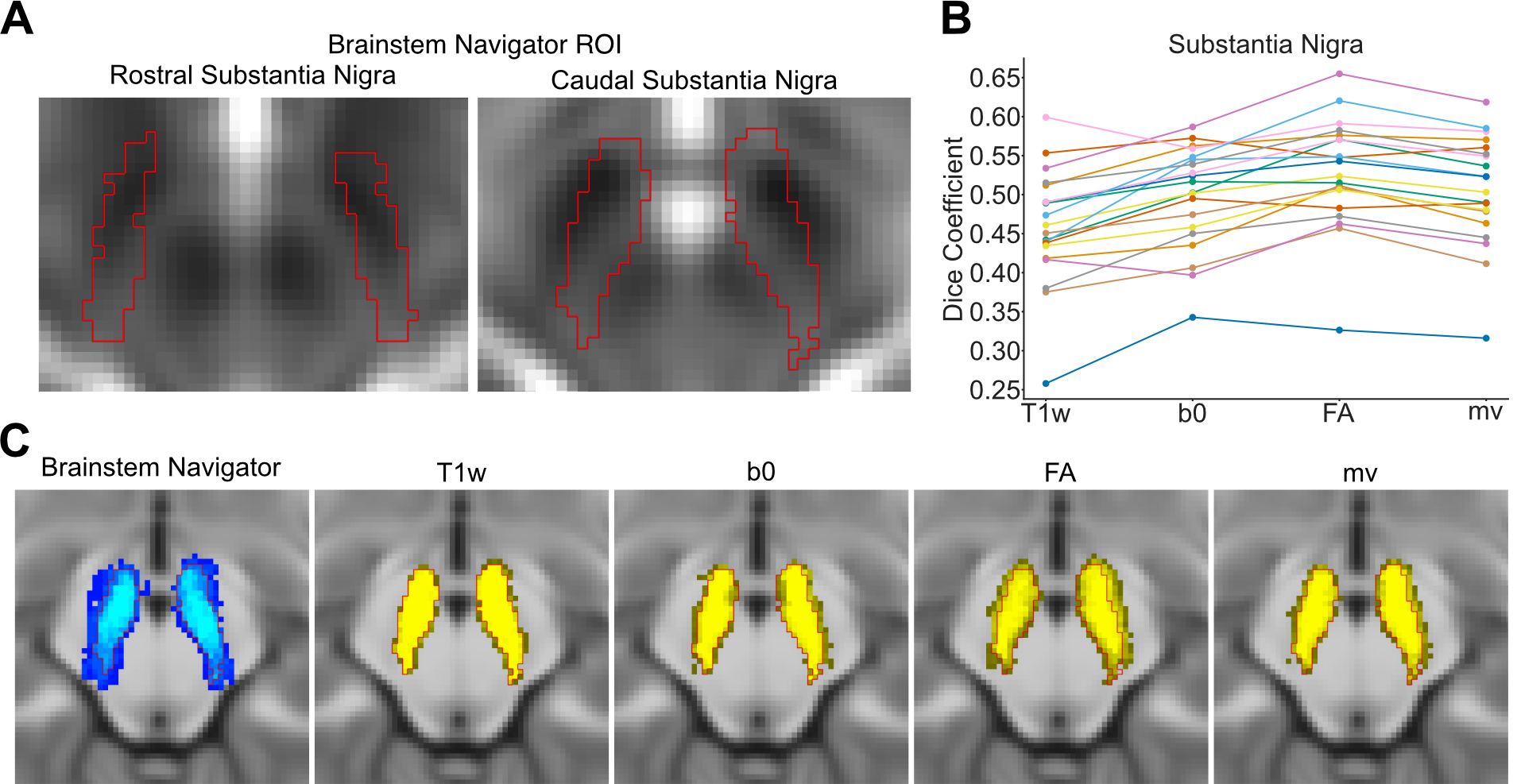

### Figure S2

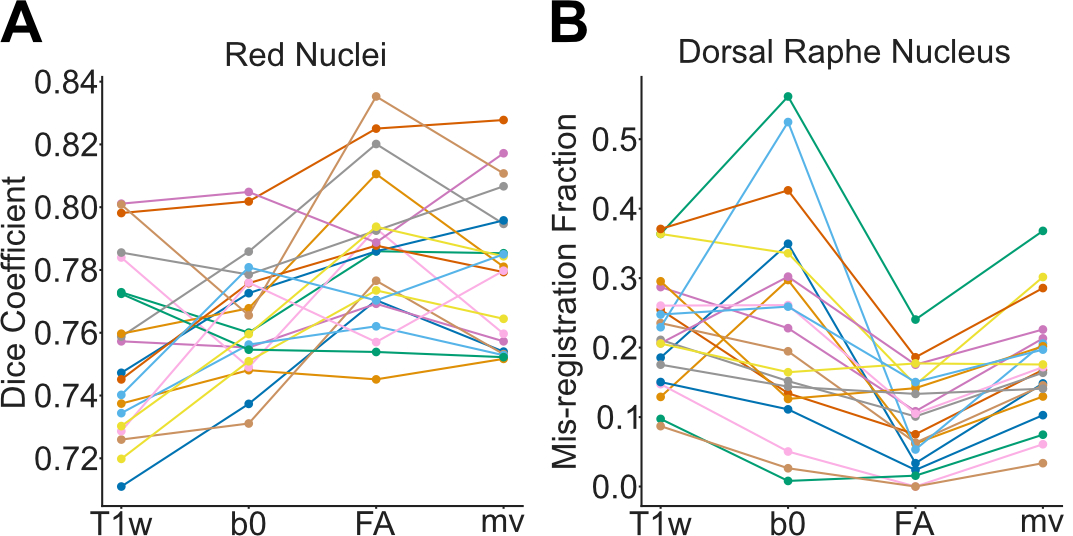

### Figure S3

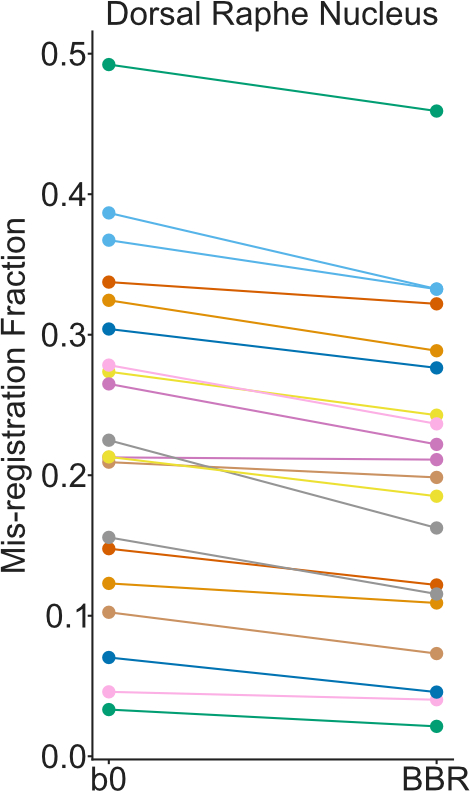
